# Arsenic Toxicity in the Drosophila Brain at Single Cell Resolution

**DOI:** 10.1101/2025.04.09.647950

**Authors:** Anurag Chaturvedi, Vijay Shankar, Bibhu Simkhada, Rachel A. Lyman, Patrick Freymuth, Elisabeth Howansky, Katelynne M. Collins, Trudy F. C. Mackay, Robert R. H. Anholt

## Abstract

Arsenic is an ubiquitous environmental toxicant with harmful physiological effects, including neurotoxicity. Modulation of arsenic-induced gene expression in the brain cannot be readily studied in human subjects. However, Drosophila allows quantification of transcriptional responses to neurotoxins at single cell resolution across the entire brain in a single analysis. We exposed *Drosophila melanogaster* to a chronic dose of NaAsO that does not cause rapid lethality and measured survival and negative geotaxis as a proxy of sensorimotor integration. Females survive longer than males but show earlier physiological impairment in climbing ability. Single-nuclei RNA sequencing showed widespread sex-antagonistic transcriptional responses with modulation of gene expression in females biased toward neuronal cell populations and in males toward glial cells. However, differentially expressed genes implicated similar biological pathways. Evolutionary conservation of fundamental processes of the nervous system enabled us to translate arsenic-induced changes in transcript abundances from the Drosophila model to orthologous human neurogenetic networks.

## Introduction

Arsenic is one of the most hazardous environmental toxicants, affecting over 140 million people globally, with computational estimates suggesting this number may exceed 200 million across more than 70 countries^1^. Arsenic toxicity is linked to diverse adverse health outcomes ^2–4^, including skin disorders ^2,3^, impaired vision ^5^, neurological damage ^6–9^, endocrine disruption ^3^, respiratory illness ^3,10,11^, and gastrointestinal distress ^4^. Biotransformation of arsenic into methylated derivatives leads to oxidative stress, which can result in genome instability and give rise to a variety of cancers ^12–14^. The accumulation of methylated derivatives also increases risk for cardiovascular disease ^15,16^. Arsenic metabolites can cross the placenta and the blood brain barrier to exert both prenatal and postnatal neurotoxic effects ^17,18^. These effects are accompanied by epigenetic modifications, which can persist trans-generationally ^19–21^.

Whereas most studies on model organisms use lethality as an endpoint, in human populations sublethal effects due to chronic exposure are more relevant, yet poorly studied. *Drosophila melanogaster* is a powerful model for toxicogenomic studies ^22,23^, since the genetic background and exposure can be controlled precisely and transcriptional responses to neurotoxins can be quantified at single cell resolution across the entire brain in a single analysis. The translational potential of the Drosophila model for arsenic neurotoxicity is underscored by well-established effects of arsenic on mitochondrial function ^18,24,25^ and oxidative stress-induced DNA damage ^13,23^, processes that are evolutionarily conserved across phyla. Whereas arsenic and its derivatives, especially meta-arsenite (NaAsO_2_), are recognized as potent neurotoxins ^5,6,8,9^, their effects on the modulation of gene expression across the human brain cannot be readily investigated. Here, we subjected a standard laboratory strain of *D. melanogaster* to a chronic dose of NaAsO and measured lifespan and climbing ability. We then performed single-nuclei RNA sequencing on male and female brains (Fig. 1). We found that the transcriptional response to arsenic neurotoxicity is characterized by widespread sex-specific and sex-antagonistic modulation of gene expression.

**Fig. 1.**
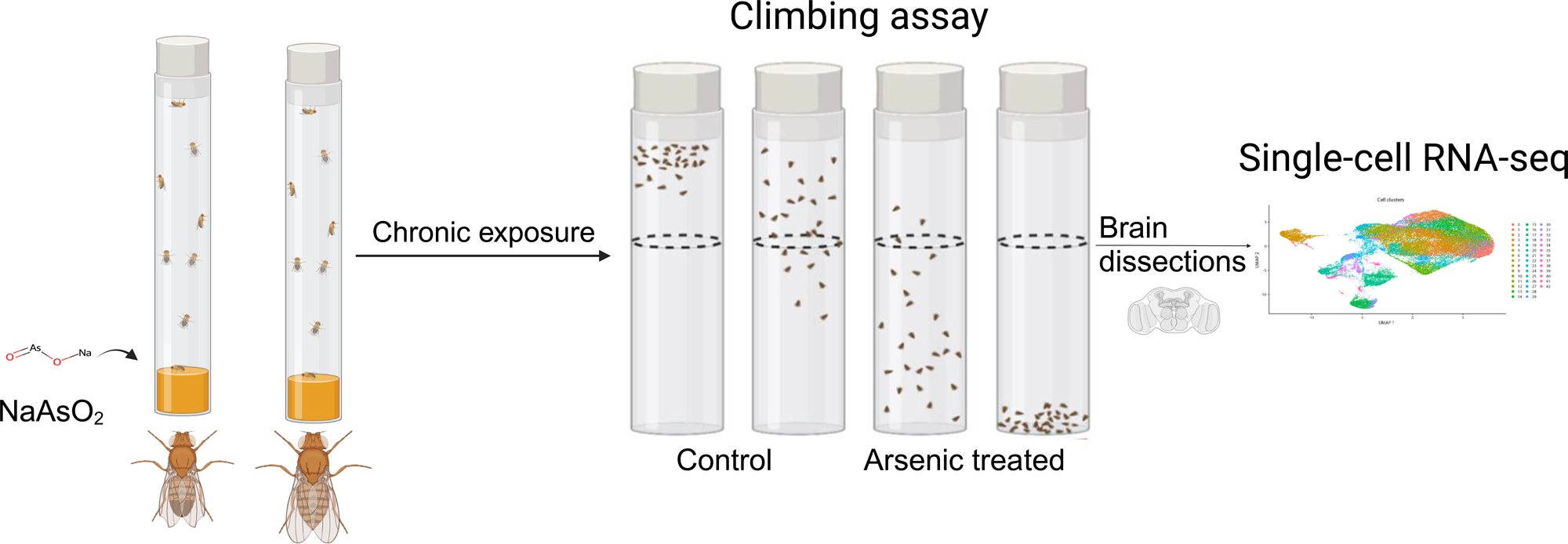
Diagram of the experimental design. Male and female flies aged between 3 to 5 days were exposed to 250 µM NaAsO_2_, followed by a climbing assay to assess locomotor activity. Single-nuclei RNA-seq analysis was then performed on dissected brains to identify molecular signatures linked to neurotoxicity.

## Results

### Chronic arsenic exposure reduces lifespan and climbing ability

We generated survival curves using a range of NaAsO concentrations and identified 250 µM as the highest concentration that showed no mortality after 24 hours of exposure in the *w*^1118^ Canton S (B) (CSB) laboratory strain (Fig. 2). Prolonged exposure to this concentration resulted in 50% mortality by day 15 (Fig. 3A, Supplementary Tabel S1) with significant effects of treatment (*P* < 0.0001) and sex × treatment (*P* < 0.0004) (Supplementary Table S2). To further assess the phenotypic effects of NaAsO exposure, we performed a climbing assay to quantify locomotor activity driven by negative geotaxis. We observed a sexually dimorphic physiological response to chronic NaAsO exposure, with females exhibiting a significant reduction in climbing ability as early as day 1, while males showed a decline in performance only after 6 days (Fig. 3B, Supplementary Tables S2 and S3). Thus, chronic exposure to concentrations of NaAsO below acute lethal levels can induce neurotoxicity that leads to adverse and sex-specific effects on survival and locomotor function.

**Fig. 2.**
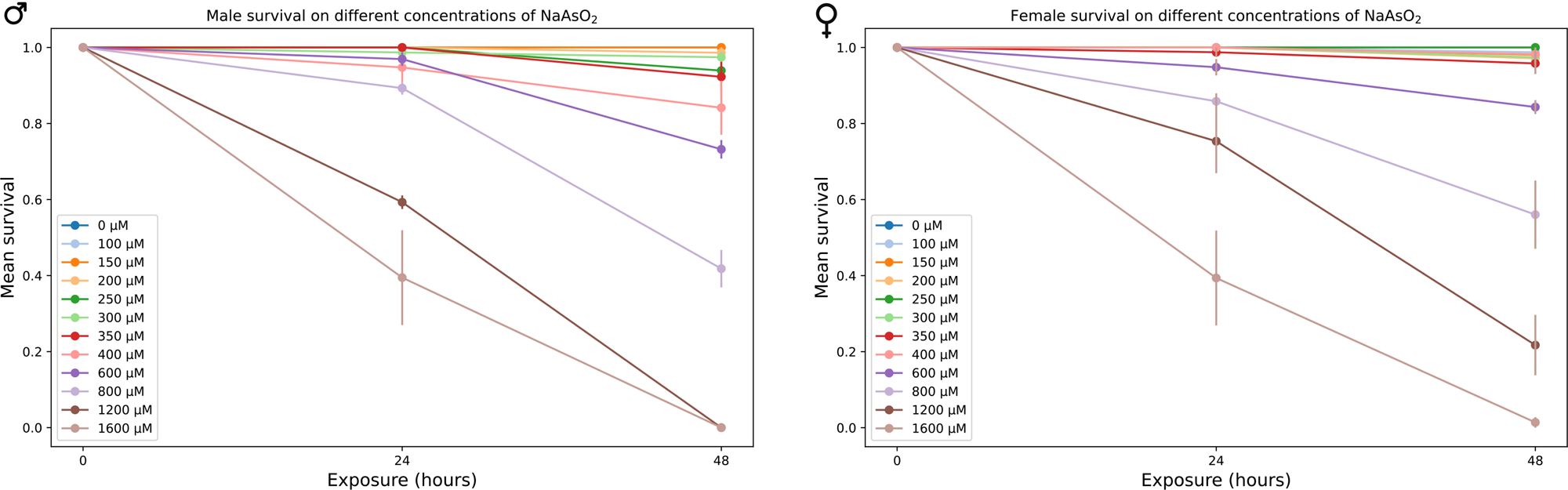
Mean survival of male and female CSB flies exposed to varying concentrations of NaAsO_2_, after 24 and 48 hrs.

**Fig. 3.**
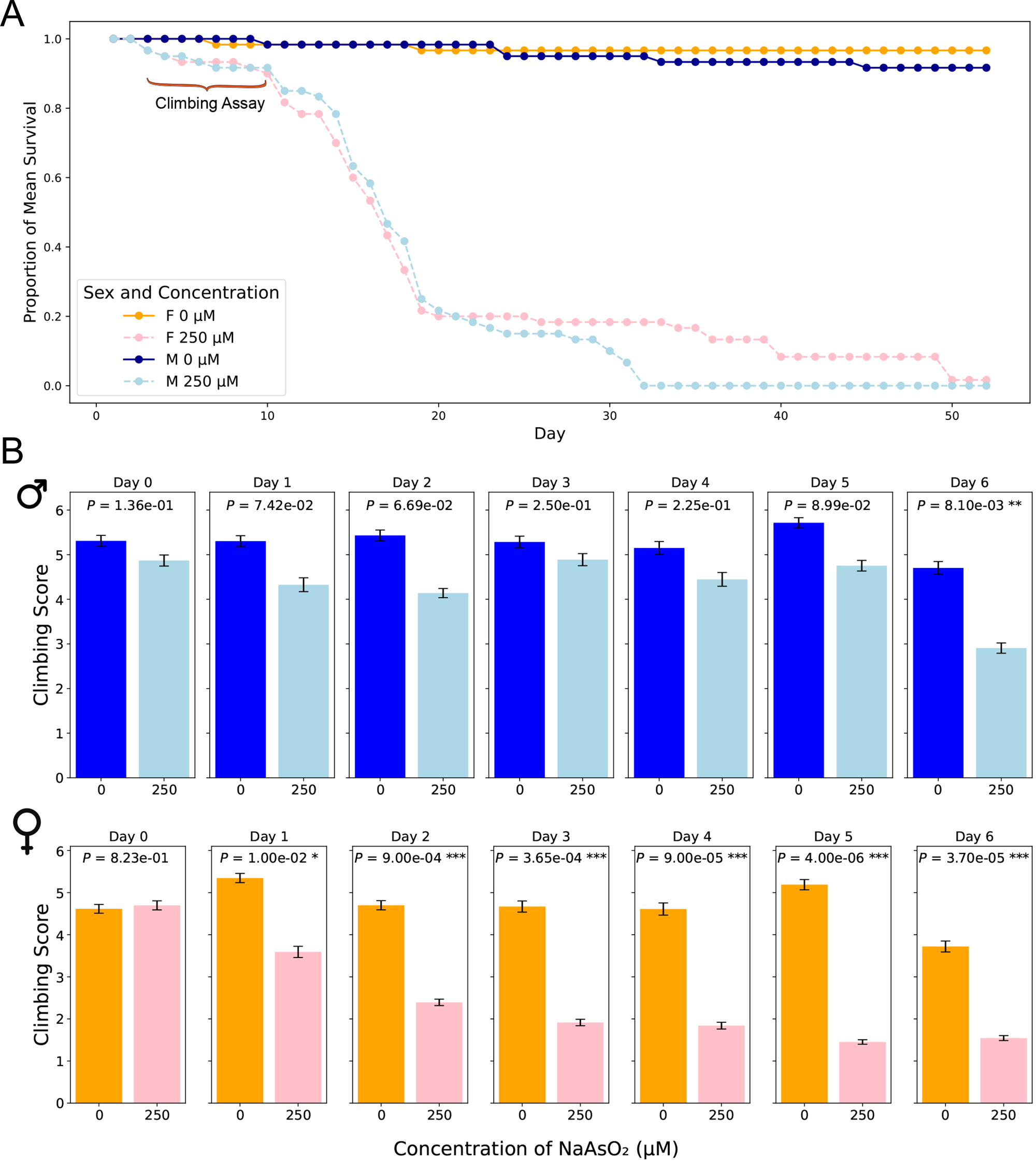
Lifespan and locomotion upon chronic NaAsO_2_ exposure. (**A**) The impact of NaAsO exposure on survival of male and female *D. melanogaster* exposed to control medium or medium supplemented with 250 µM NaAsO. (**B**) The impact of NaAsO exposure on climbing ability.

### Sexually antagonistic transcriptional responses to chronic NaAsOD exposure in the Drosophila brain at single cell resolution

We performed snRNA-seq analysis for pools of brains from CSB males and females collected after 6 days of exposure to 250 µM NaAsO and from control flies of the same age not exposed to NaAsO. We profiled whole brain transcriptomes of 68,709 single nuclei (Supplementary Table S4) and identified 35 cell clusters representing 20 major neural and glial cell types, including those from the optic lobe, mushroom body, and blood-brain barrier (Fig. 4, Supplementary Fig. S1, Supplementary Table S5). Our differential expression analysis revealed changes in 767 genes, with 53.7% (412 genes) with altered transcript abundances in both males and females, 18.6% (143 genes) unique to females, and 27.6% (212 genes) unique to males (Supplementary Fig. S2 and S3, Supplementary Table S6). The differentially expressed genes were biased toward neurons in females and glia in males (Fig. 5).

**Fig. 4.**
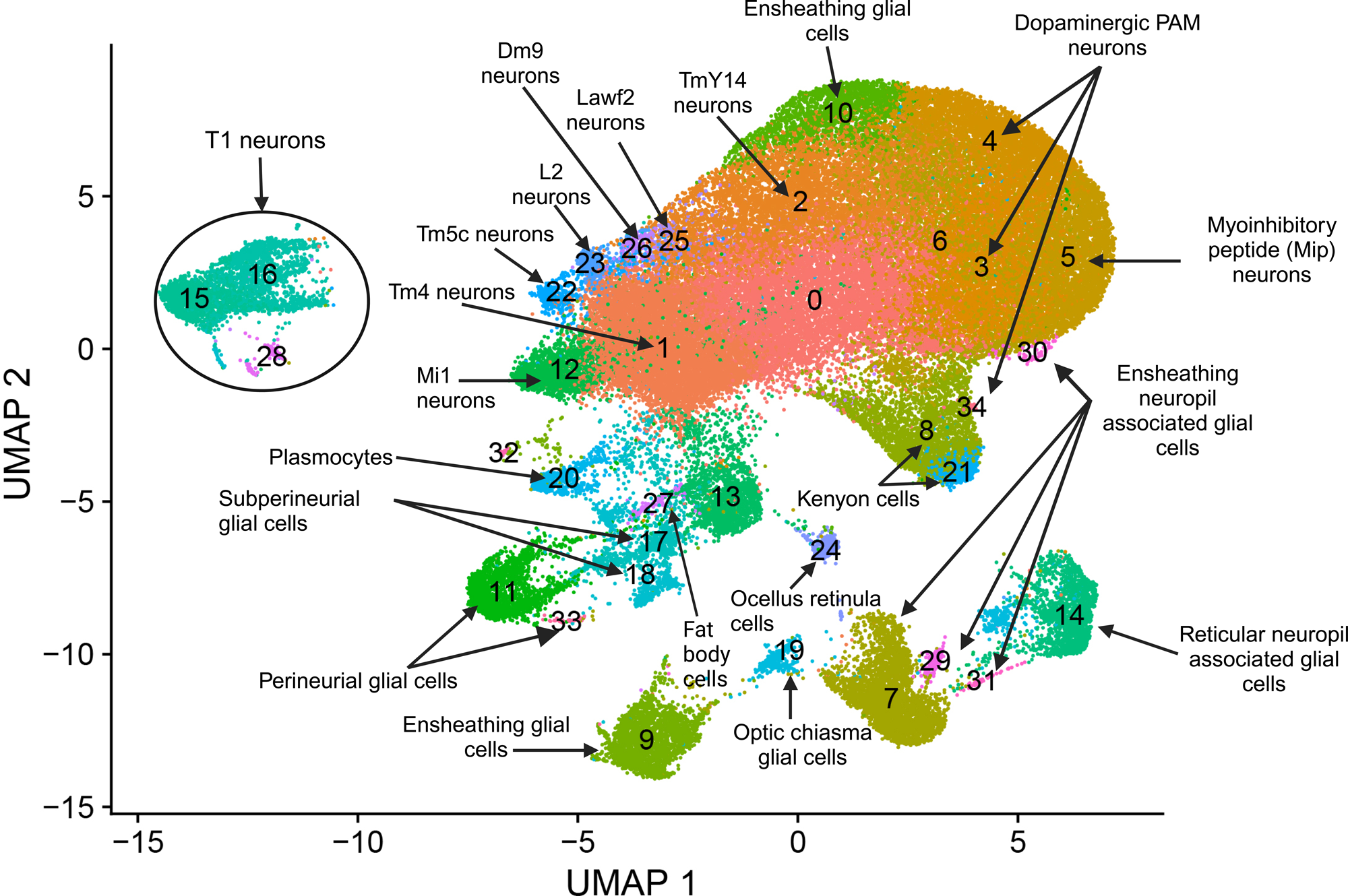
snRNA-seq analysis of brains from flies exposed to chronic NaAsO_2_ exposure. Cells were clustered according to their gene expression patterns using the unsupervised Shared Nearest Neighbor (SNN) clustering algorithm. Each dot represents an individual cell, with colors indicating the cluster to which the cell belongs. Cell type identification for each cluster was performed by annotating the top 20 marker genes. These markers were determined based on the most frequently occurring cell type identities using expression scores obtained from the BGEE database ^58^.

**Fig. 5.**
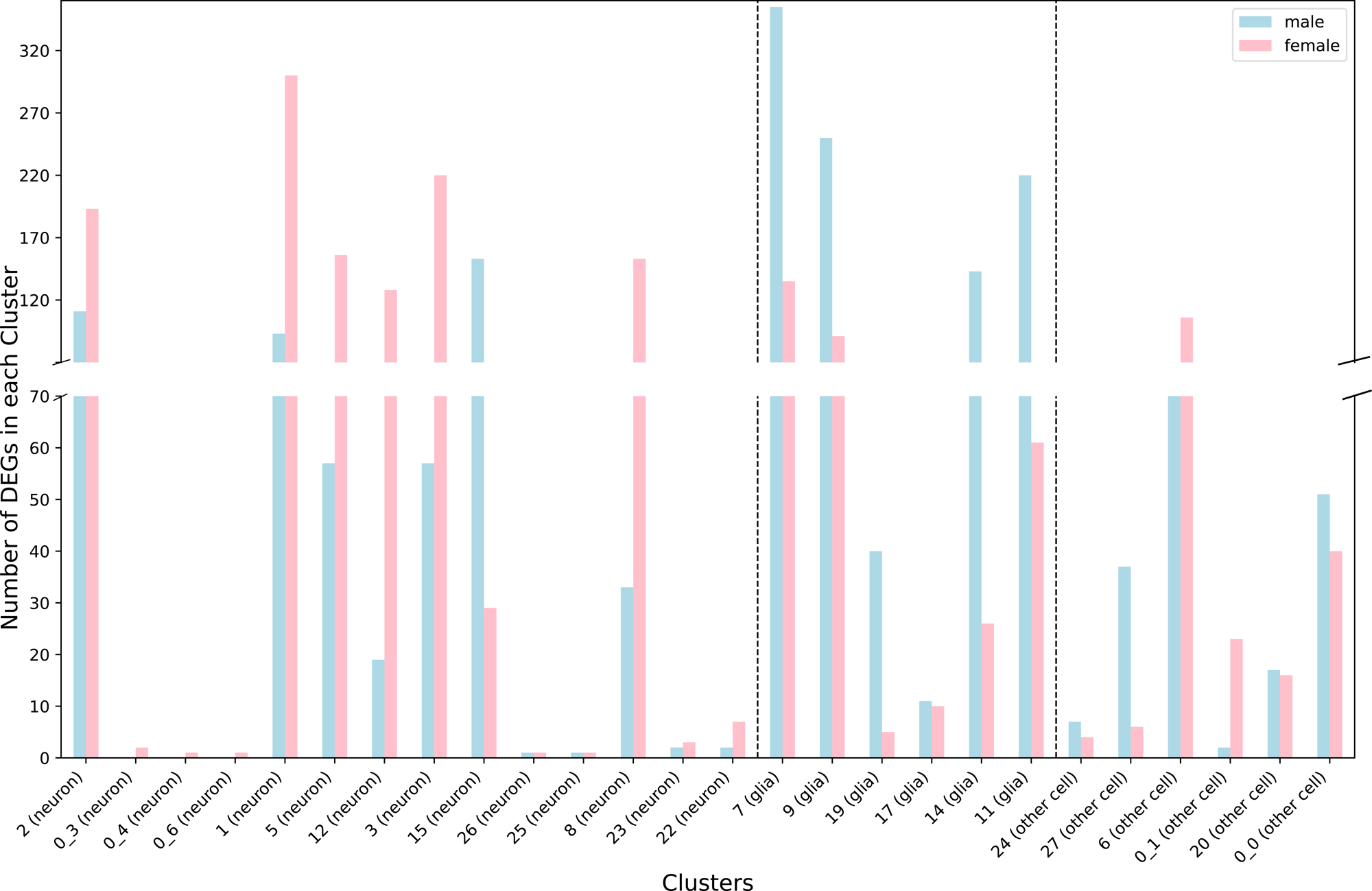
Distribution of differentially expressed genes between neurons and glia across cell clusters for males (blue) and females (pink) separately.

We observed that most gene expression differences between control and NaAsO-treated flies were sexually antagonistic, with 94.7% (390 of the 412 shared genes) of differentially expressed genes in females showing downregulation, while 82.7% (341 of the 412 shared genes) in males exhibited upregulation across cell types (Fig. 6, Supplementary Fig. S4). Cell type-specific sex-specific expression was also evident, with 10 genes in females and 26 genes in males exhibiting expression profiles in multiple distinct cell types (Supplementary Table S7).

**Fig. 6.**
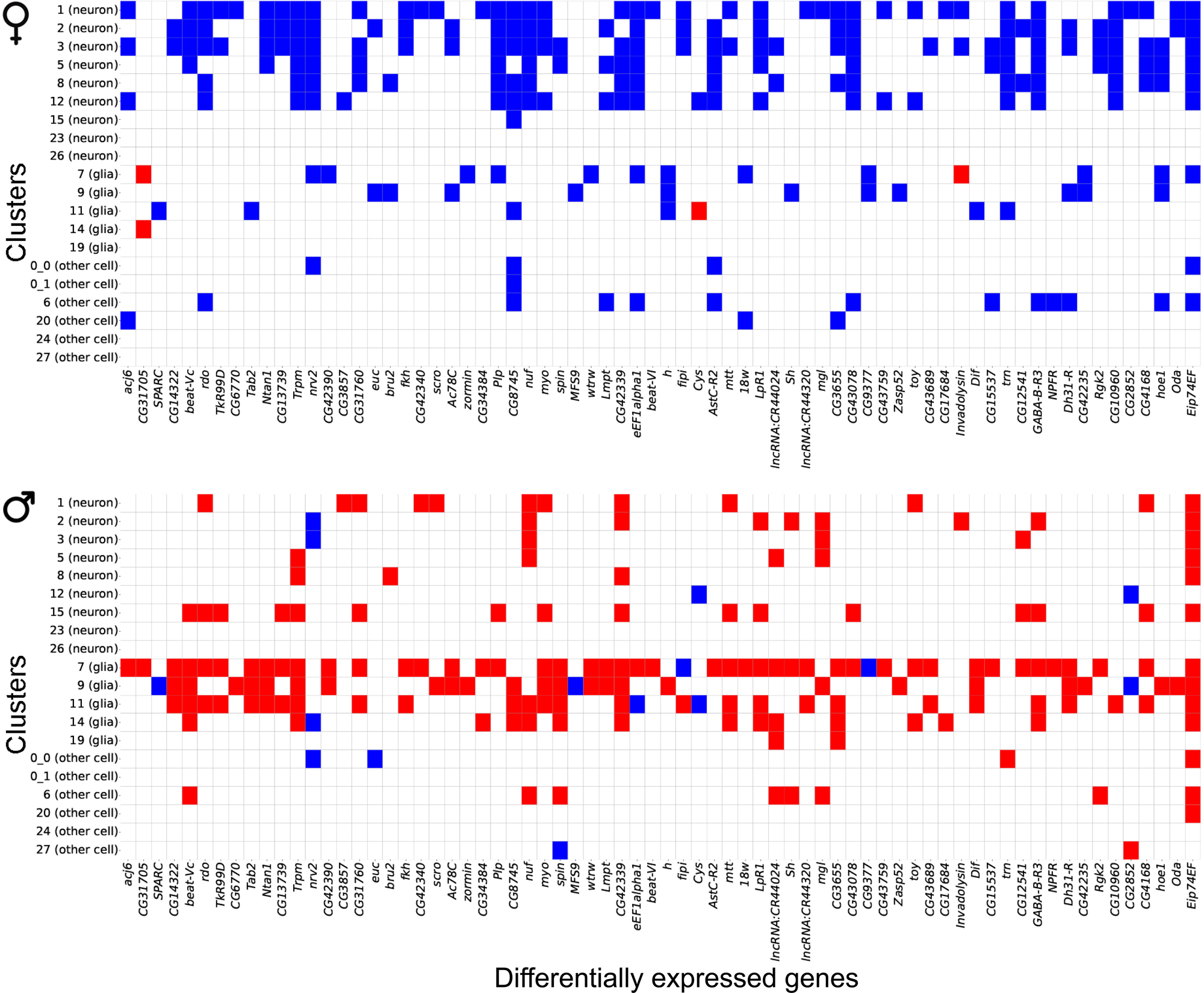
Gene expression patterns between males and females following chronic NaAsOD exposure. Heatmap showing antagonistic gene expression patterns between males and females for 70 representative genes out of 412 shared DEGs across neuronal and glial cell types.

Further analysis of differentially expressed gene profiles revealed dysregulation of genes associated with all major neurotransmitter receptor and transport systems, including cholinergic, dopaminergic, GABAergic, and glutamatergic systems (Fig. 7). These NaAsO-induced changes in gene expression involved receptor subunits and transporters critical for neurotransmitter synthesis, release, reuptake, and signaling (Fig. 7). We observed downregulation of both inhibitory and excitatory neurotransmitter-associated genes in both sexes. For example, *Vglut* (vesicular glutamate transporter), *Vmat* (vesicular monoamine transporter), and *Gat* (GABA transporter) were downregulated in both sexes, albeit in different cell types. Thus, chronic exposure to NaAsO leads to widespread disruption in synaptic communication.

**Fig. 7.**
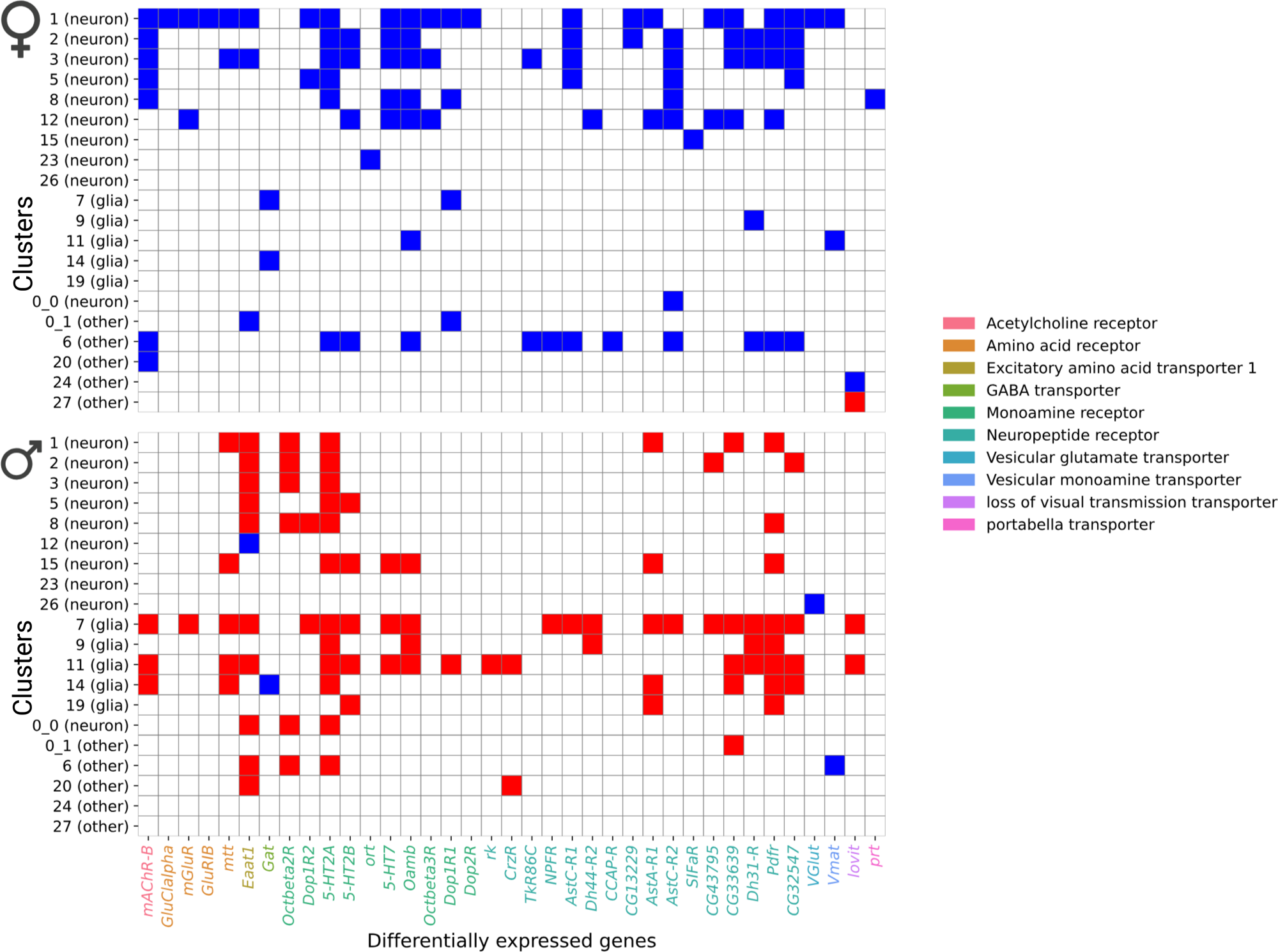
Heatmap showing differentially expressed genes in neurotransmitter receptor and transporter categories. Blue shows down-regulated genes and red shows up-regulated genes.

### Differential expression of genes associated with detoxification

The Drosophila fat body, functionally analogous to mammalian adipose tissue and liver, plays a central role in heavy metal detoxification ^26^. We recovered head fat body tissue in our dissected brain samples. Metallothionein (*Mtna*) and Turandot (*Tot*) genes are critical for responding to heavy metal toxicity ^27^. We observed the highest upregulation of gene expression for *Mtna* in the fat body (cluster 27), with a 42-fold increase in males and a 43-fold increase in females. Conversely, we found significant downregulation for *TotA* (70-fold) and *TotC* (48-fold) in the fat body of males (Supplementary Table S6).

Functional enrichment analysis of upregulated genes in males revealed pathways related to the metabolism of xenobiotics by cytochrome P450 and members of the Glutathione S-transferase (GST) gene family (Supplementary Table S8). *Cyp28d1* features prominently in both males and females, where it is upregulated in Cluster 7, associated with ensheathing neuropil-associated glial cells. In males, *Cyp28d1* is also upregulated in optic chiasma glial cells (Cluster 19) and downregulated in fat body cells (Cluster 27). Expression of *Cyp28a5* is altered in Clusters 7 and 9 in males. Additional members of the cytochrome P450 family (*Cyp4g15*, *Cyp6w1* and *Cyp311a1*) also show upregulation in male glial cells (Cluster 17). In females, expression of *GstD1* is downregulated in Kenyon cells (Cluster 8), while it is upregulated in fat body cells in males (Cluster 27). Similarly, *GstE6* and *GstE9* are both upregulated in fat body cells in males, whereas *GstD9* is downregulated in ensheathing glial cells (Cluster 9).

Members of the UDP-glucuronosyl transferase (UGT) gene family also show differential changes in brain transcript abundances in flies exposed to NaAsO (Supplementary Table S6). In males, expression of *Ugt49C1* is increased in fat body cells (Cluster 27) and ensheathing glial cells (Cluster 9) and perineurial glial cells (Cluster 11). Conversely, expression of *Ugt35B1* is reduced in males in ensheathing glial cells (Cluster 9) and ensheathing neuropil-associated glial cells (Cluster 7). In females, *Ugt35E2* is downregulated in ensheathing glial cells (Cluster 9), and *Ugt35B1* is reduced in ensheathing neuropil-associated glial cells (Cluster 7). These patterns indicate largely cell-and sex-specific roles of NaAsO_2_-induced Cytochrome P450s, GSTs, and UGTs in detoxification.

Complementing these detoxification enzymes, transcripts encoding *MsrA* (*Methionine sulfoxide reductase A*) and *Sod3* (*Superoxide dismutase 3*), which neutralize superoxide radicals generated during oxidative stress ^28–30^, are upregulated in males, particularly in fat body cells (Cluster 27), with up-regulation in ensheathing glial cells (Cluster 9), perineurial glial cells (Cluster 11), and ensheathing neuropil-associated glial cells (Cluster 7). Expression of *Sestrin* (*Sesn*), which plays a vital role in oxidative stress response ^31^, is downregulated in females in Myoinhibitory peptide (MIP) neurons, ensheathing neuropil-associated glial cells, and dopaminergic PAM neurons; and upregulated in males in ensheathing glial cells, reticular neuropil-associated glial cells, ensheathing neuropil-associated glial cells, and perineurial glial cells.

### Differential expression of genes associated with the blood brain barrier and ensheathing glia

The Drosophila blood-brain barrier (BBB) consists of perineurial and sub-perineurial cells, which play critical roles in maintaining neural homeostasis ^34^. In perineurial cells (Cluster 11), downregulated genes were enriched in the Notch signaling pathway, previously shown to be essential for maintaining the BBB integrity in subperineurial glia by regulating cell proliferation, differentiation, and the expression of tight junction proteins ^35^. In males, upregulated genes in these cells were predominantly enriched in the heterotrimeric G-protein signaling pathways, specifically Gα_i_ – and Gα_s_-mediated signaling pathways (Supplementary Table S8).

BBB cells establish contacts with ensheathing neuropil-associated glial cells ^36^. In ensheathing glial cells, NaAsO_2_ exposure elicited distinct sex-specific responses. In females, downregulated genes were enriched in pathways associated with the transport of small molecules, SLC-mediated transmembrane transport, and GABA-B receptor activation. In contrast, upregulated genes in males were enriched in heterotrimeric G-protein signaling pathways and the endothelin signaling pathway. Notably, components of GPCR signaling, Notch signaling, MAPK signaling and adenylate cyclase inhibitory pathways were enriched in ensheathing glial cells of both sexes (Supplementary Table S8). However, these pathways exhibited antagonistic responses in differential gene expression patterns between males and females (Supplementary Table S6).

### Differential expression of genes associated with vision and locomotion

Since there were profound effects of chronic NaAsO_2_ exposure on negative geotaxis (Fig. 3), we examined the extent to which genes associated with sensorimotor integration and locomotion showed NaAsO_2_-induced changes in gene expression in the brain. We identified 102 genes in 10 male cell clusters (0_0, 1, 2, 6, 7, 8, 9, 11, 14, 15) and 96 genes in nine female clusters (1, 2, 3, 5, 7, 8, 9, 11, 22), enriched for locomotion-related functions in both neuronal and glial cell types (Supplementary Table S8). We observed the most significant responses to NaAsO_2_ exposure in genes involved in memory, vision, and motion. Notably, the *Rhodopsin 2* (*Rh2*) gene exhibited the largest downregulation, particularly in the ocellus retinula cells, with an average 25-fold and 13-fold decrease in males and females, respectively (Supplementary Table S6). Genes involved in these processes were organized in co-expression networks, highlighting multiple neuronal cell types in the optic lobes implicated in vision and locomotion, including Tm4, TmY14, T1, Tm5c, L2, Lawf2, Dm9, LC17, T3, Tm9 and Mi1 neurons ^37^ (Fig. 8).

**Fig. 8.**
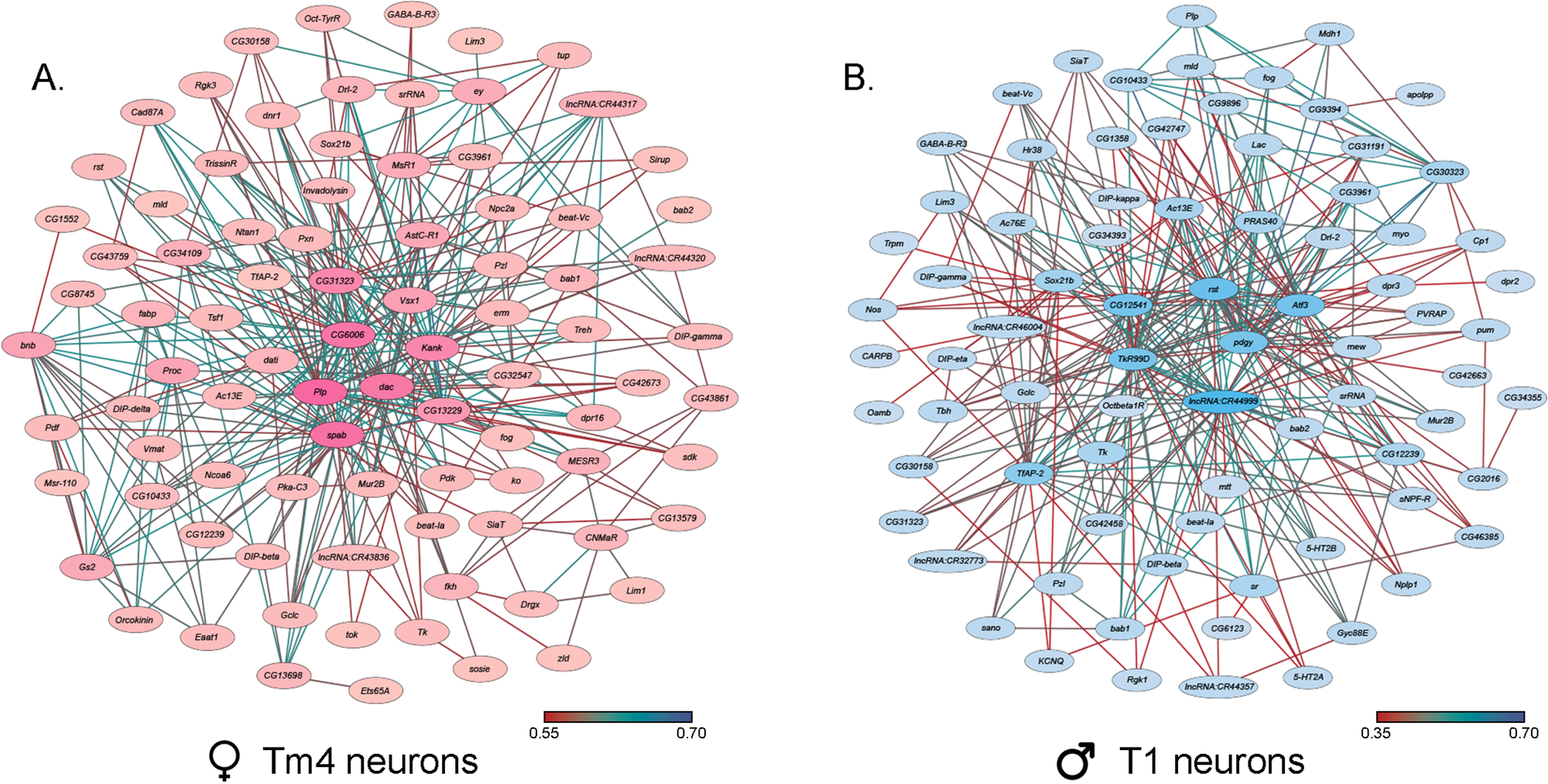
Co-expression networks of neurons in the optic lobes that exhibit the largest responses to NaAsO_2_ exposure. (**A**) Co-expression network of Tm4 neurons in female cluster 1. (**B**) Co-expression network of T1 neurons in male cluster 15. The darker colors in both coexpression networks represent a higher number of connections.

Expression of *Arrestin 2* (*Arr2*), associated with phototransduction, was downregulated in photoreceptor neurons in females and multiple male neuron types, including myoinhibitory peptide (MIP) neurons, which regulate reproductive behaviors ^38^, and dopaminergic PAM neurons, critical for reward processing and learning ^39–41^ (Supplementary Table S6). In males, expression of the *Ndae1* (Na^+^/H^+^ antiporter) gene, which can mediate clearance of toxic arsenic anions ^42^, is upregulated in optic chiasma glial cells, reticular neuropil-associated glial cells, and Tm4 neurons. In females, *Ndae1* expression is downregulated in Kenyon cells, MIP neurons, and dopaminergic PAM neurons. Additionally, differentially expressed genes in Kenyon cells, which mediate experience-dependent modification of behavior, showed sex-specific responses in G protein-coupled receptor signaling pathways. These results emphasize the wide-ranging effects of NaAsO_2_ exposure on neuronal function, with distinct impacts on cells associated with sensory processing and motor coordination.

### Global protein-protein interaction networks

Although the arsenic-induced changes in transcript abundances differ in a sex-specific manner, the overarching biological processes disrupted by NaAsO_2_ exposure are shared between males and females (Supplementary Table S9). For example, in pathways related to neurotransmitter signaling and synaptic transmission, genes involved in receptor activation and signal propagation were upregulated in males (Fig. 7). In contrast, genes associated with neurotransmitter synthesis and vesicle transport were predominantly downregulated in females. Similarly, G protein-coupled receptor (GPCR) signaling and calcium signaling in oxidative stress response pathways were enriched in both sexes but were modulated through different sets of genes.

We analyzed global protein-protein interaction networks of differentially expressed gene products across all cell clusters for both males and females to identify biological functions impacted by NaAsO_2_ exposure. Among differentially expressed genes in the Drosophila brain 72.35% had human orthologs. Despite pronounced sexual dimorphism and antagonism in gene expression responses to arsenic, arsenic-induced neurotoxicity affected the same pathways in both males and females but involved distinct components within these pathways. We constructed global protein-protein interaction networks in which we computationally recruited known physically interacting partners with the products of the differentially expressed genes (Fig. 9).

**Fig. 9.**
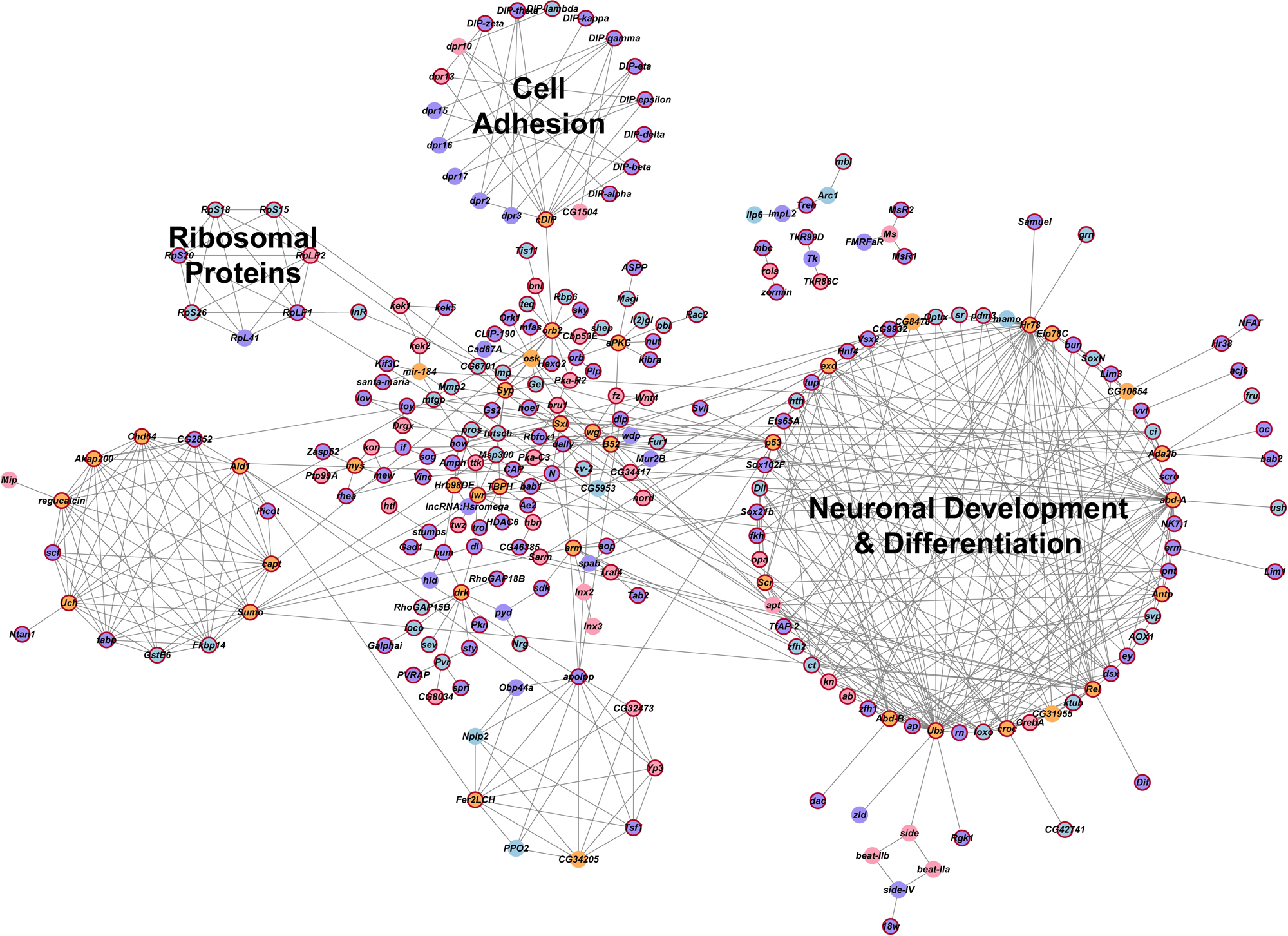
Global gene interaction network. The protein-protein interaction network was constructed within Cytoscape, based on interactions among genes that were differentially expressed (Benjamini-Hochberg adjusted *P*-value ≤ 0.05) across all clusters in the male and female datasets. Crimson red borders indicate genes with human orthologs. Circles representing genes are color-coded: purple indicates genes differentially expressed in both males and females, blue indicates genes differentially expressed only in males, pink indicates genes differentially expressed only in females and orange indicates inferred genes that were computationally recruited from the FlyBase Interaction Database. Annotations of these groups represent the processes that are enriched for the genes within these groups. A Benjamini-Hochberg ^67^ adjusted *P*-value ≤ 0.05 was considered a significant enrichment in the statistical overrepresentation tests.

The global network consists of 258 nodes and reveals distinct subnetworks, including a large subnetwork associated with neuronal development and differentiation, a subnetwork of cell adhesion proteins recruited through cDIP, and a cluster of ribosomal proteins (Fig. 9). Based on evolutionarily conserved orthology, we constructed a human network corresponding to the Drosophila global protein-protein interaction network (Fig. 10A and B). Comparisons between these networks reveal common edges that show a conserved core structure of protein-protein interactions associated with the transcriptional response to arsenic neurotoxicity (Fig. 10C). These include ribosomal protein interactions, signal transduction pathways, and developmental processes (Fig. 10C). Statistical analysis shows that the likelihood of obtaining this orthologous network architecture due to randomness is *P* < 0.0001 (Supplementary Fig. S5).

**Fig. 10.**
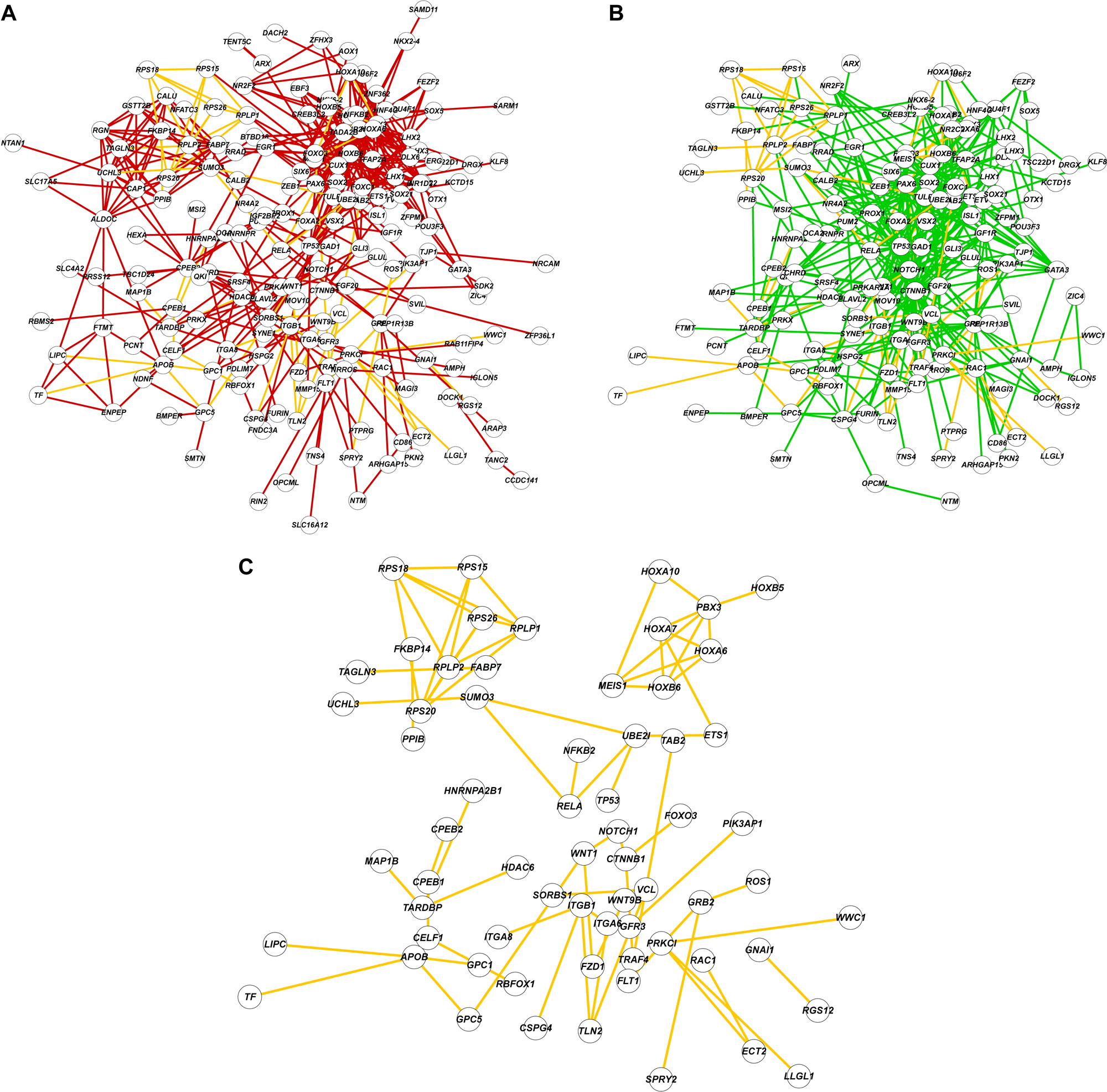
Comparisons of protein-protein interaction networks. (**A**). The protein-protein interaction network constructed with Drosophila genes and FlyBase physical interaction database. Nodes are labeled with human orthologs of Drosophila genes. (B). The protein-protein interaction network constructed with human orthologs and interactions from String-DB database. (C). A subnetwork composed of shared edges (yellow) between A and B.

## Discussion

Exposure to NaAsO_2_ at a concentration that does not result in immediate lethality affects Drosophila locomotor ability sooner in females than in males. The transcriptional response in the brain after six days of chronic exposure to NaAsO_2_ is widespread and different between males and females, yet gene ontology analysis suggests that these sexually dimorphic responses converge through different mechanisms on the same biological processes. The transcriptional responses to arsenic exposure are markedly sex antagonistic with responses in females biased toward neurons and in males toward glia. The mechanisms that underlie sex antagonistic modulation of arsenic-induced gene expression are not known, but may arise from differences in chromatin accessibility, differences in chromosomal conformations due to the presence of different sex chromosomes affecting intrachromosomal interactions, as well as antagonistic regulation of the *doublesex* (*dsx*) gene, which shows NaAsO_2_-induced altered transcript abundance and is part of the sex determination pathway ^43^.

Chronic exposure to NaAsO_2_ leads to differential expression of more than 5% of the Drosophila genome, indicating a highly polygenic response that is likely to affect multiple organismal phenotypes. We used innate negative geotaxis as a proxy phenotype for NaAsO_2_ neurotoxicity as it can readily be quantified using a climbing assay. It is, however, likely that phototaxis is also affected by NaAsO_2_ exposure, based on the prominent changes in transcript abundances in cell clusters associated with vision. In addition, gene ontology enrichment analysis implicates possible additional physiological, developmental and behavioral phenotypes that may be affected by NaAsO_2_, including locomotion, mating and courtship, feeding, sleep and circadian rhythm as well as neurodevelopment and nervous system function, including sensory perception. Locomotion and circadian rhythm have previously been implicated in studies on arsenic toxicity in Drosophila ^10^.

Differential regulation of genes associated with detoxification processes and response to oxidative stress features prominently in the transcriptional response to chronic NaAsO_2_ exposure, most notably in males in glial cell populations, in concordance with their greater resilience to NaAsO_2_ than females as reflected in their climbing ability. In addition, many genes associated with locomotion were differentially expressed upon chronic exposure to NaAsO_2_, including the *X*-chromosome-linked genes *spin, sr, tim, zfh1, dac, ey, rhea, robo2, and Sh,* also in line with the adverse effect of NaAsO_2_ on climbing ability.

Over 60% of differentially expressed genes occur in multiple cell clusters in both males and females. In females, the top 10 genes (*Orcokinin*, *Pdf*, *Ms*, *fabp*, *CG12239*, *Nplp1*, *lncRNA:CR40469*, *CG46385*, *CG8745*, and *ATP8B*) were differentially expressed across multiple clusters. In males, *srRNA*, *TotC*, *CG12239*, *Eip74EF*, *Pzl*, l*ncRNA:CR46004*, *apolpp*, *5-HT2A*, T*otA*, *sr*, *Obp44a*, *Eaat1*, *puc*, *MtnA*, and *CG34393* showed widespread differential expression (Supplementary Table S10).

*Orcokinin* and *Nplp1* are differentially expressed in females in 20 and 18 clusters, respectively, and have been implicated in sleep, locomotion, and feeding ^41,42^. In males, *Eip74EF*, a target of the ecdysone signaling pathway, associated with metamorphosis, apoptosis, immune response, and stress ^44–47^ is differentially regulated in 13 clusters. The gene *puc* (*puckered*), a negative regulator of JNK signaling, that contributes to apoptosis, stress responses, and immune function ^48^ is differentially expressed in 10 cell clusters in response to chronic NaAsO_2_ exposure.

Additionally, transcript abundance of the 5-HT2A serotonin receptor, which modulates neural signaling related to feeding and locomotion ^49^, is altered in 12 cell clusters. These examples underscore the widespread pleiotropic effects on diverse physiological and behavioral phenotypes that are likely to result from arsenic exposure.

Previous studies have shown extensive genetic variation in susceptibility to cadmium and lead ^50^ in an advanced intercross population derived from the inbred lines of the Drosophila Genetic Reference Panel (DGRP) ^51,52^. Variation in sexual dimorphism in susceptibility to 4-methylimidazole has also been documented in the DGRP ^53^. It is, however, likely that the widespread changes in gene regulation across multiple cell populations in the brain and the sexual dimorphism and sexual antagonism that are characteristic of the neurotranscriptional response to NaAsO_2_ will also be representative for transcriptional responses to other neurotoxins. Furthermore, we analyzed concordance between Drosophila and human protein-protein interaction networks and found statistically significant conservation of functional relationships indicating our ability to infer neurotoxic mechanisms in the human brain from the Drosophila model.

## Methods

### Drosophila stock

Canton S (B) flies (CSB) were reared on standard cornmeal-yeast-molasses-agar medium at 25°C under a 12:12 h light:dark cycle and 50% humidity. Five males and five females were placed in each vial to mate for 2 days before being cleared to prevent overcrowding. Progeny were collected after eclosion by brief exposure to CO_2_ anesthesia and aged for 3–5 days prior to experimentation.

### Effective range determination

Sodium (meta) arsenite (NaAsO) (CAS Number: 7784-46-5) was purchased from Sigma-Aldrich, and a range-finding assessment was conducted using CSB flies to determine the concentration of NaAsO that supported survival after 24 and 48 hours of exposure for both sexes, following a modified version of a previously established protocol ^54^. Two weeks prior to exposure, controlled adult density vials were set up with five female and five male CSB flies per vial, and after 48 hours, flies were transferred to fresh vials to promote egg-laying. Adults were removed 48 hours later, and after seven days, progeny were anesthetized using CO and sorted into groups of 15–20 same-sex flies. At three to five days post-eclosion, following a minimum three-hour recovery period from anesthesia, flies were food-deprived for 20 hours in vials containing three pieces of 20 mm Whatman #1 filter paper (Cytiva Cat. #1001-020) wetted with 0.35 mL distilled water. Subsequently, flies were exposed for 24 and 48 hours to 0, 100, 150, 200, 250, 300, 350, 400, 600, 800,1200 and 1600 µM NaAsO by transferring them to vials containing three pieces of 20 mm filter paper wetted with 0.35 mL of an exposure solution comprising NaAsO, 1.5% yeast, 4% sucrose, and distilled water, with four biological replicates of 15–20 individuals per sex/concentration. Survival was recorded for each replicate and averaged across all replicates, with standard errors calculated as 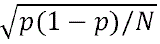, where p is the pooled proportion surviving across replicates, and N is the total number of individuals assessed.

### Climbing assay

Negative geotaxis was quantified using a climbing assay with a vertically oriented Benzer’s counter-current apparatus on 3-5 day old flies. Control flies were maintained on standard medium at 25°C under a 12:12 h light:dark cycle and 50% humidity without chemical exposure. Treated flies were maintained under the same conditions, but their medium was supplemented with 250 µM NaAsO. Flies were grouped by sex and exposure day into sets of 20 individuals, and the climbing activity of both control and treatment groups (three replicates each for each sex) was recorded daily in the morning from 9 to 11 AM over six consecutive days. A new set of flies was used on each assay day. For each replicate, flies were collected without anesthesia and allowed to acclimate for 1-2 hours. During the assay, flies were gently tapped to the bottom of the starting tube and given 15 seconds to climb into the distal tube. Flies reaching the distal tube were transferred to the next tube, and this process was repeated seven times. At the end of the assay, flies in all eight tubes were frozen, counted, and scored from 1 (indicating no successful crossings) to 8 (indicating successful crossings in all seven trials).

Climbing scores were analyzed using a mixed-effects ANOVA with the PROC MIXED procedure in SAS™ Studio v3.9 ^55^. The full model used the Type III method to test significance and was defined as follows: Y = µ + treatment (T, fixed) + Sex (S, fixed) + Day (D, fixed) + TxS, TxD, SxD, TxSxD (all fixed) + rep(TxSxD) (random) + ε. In addition, a reduced ANOVA model Y = µ + T (fixed) + rep(T) (random) + ε was used to assess climbing performance by day and sex.

### Lifespan assay

Lifespan assays were conducted on males and females separately by exposing them to either standard medium or medium supplemented with 250 µM NaAsO. To prevent overcrowding, five males and five females were placed in each vial to mate for 2 days. Progeny were collected after eclosion, and lifespan was assessed by transferring them without anesthesia to new vials with fresh medium daily and recording deaths until all NaAsO-treated flies had died. We measured 20 replicate vials per treatment, each containing 5 mL of culture medium and 3 same-sex flies. Survival was recorded as means across all replicates. The full model used the Type III method to test lifespan significance and was defined as follows: Y = µ + treatment (T, fixed) + Sex (S, fixed) + TxS (fixed) + rep(TxS) (random) + ε were analyzed using a mixed-effects ANOVA with the PROC MIXED procedure in SAS™ Studio v3.9 ^55^.

### Brain dissection and single nucleus RNA sequencing (snRNA-seq)

We anesthetized CSB flies on ice and dissected 10 brains per sex after six days of exposure to control medium or medium supplemented with 250 µM NaAsO_2_ in cold Dulbecco’s phosphate-buffered saline (PBS). Brains were flash frozen in a dry ice-ethanol bath, and stored at –80°C. We prepared the samples for snRNA-seq as described in the 10X Genomics Gene Expression (GEX) protocol (CGOOO338). 200 μl of chilled Lysis Buffer was added to each tube containing the brain samples and incubated on ice for 5 minutes. The brains were mechanically dissociated through stepwise trituration with a P200 micropipette (5 times), a 23G needle (5 times), and a 27G needle (5 times). The dissociated samples were passed through a 30 μm MACS SmartStrainer (Miltenyi Biotec B.V. & Co. KG), followed by rinsing with 800 μl of lysis buffer, and collected into 5ml tubes. Each sample was then passed through a 10 μm strainer (Celltrics, Görlitz, Germany), followed by rinsing with 1 ml PBS + 1% BSA, and collected into a new 5ml tube. A 20 μl aliquot was removed for quantitative analysis, and the remaining sample was centrifuged at 500 rcf for 5 minutes at 4°C. After centrifugation, the supernatant was removed, and the pellet was resuspended in a chilled diluted nuclei buffer ^56^. Single nuclei were counted using a hemocytometer with trypan blue exclusion and proceeded to GEM generation using the microfluidics Chromium Controller (10X Genomics, Pleasanton, CA). Libraries were prepared following the 10X Genomics v3.1 protocols. Fragment sizes were determined using Agilent Tapestation High Sensitivity D5000 for amplified cDNA (Agilent Technologies Inc., Santa Clara, CA) and High Sensitivity D1000 ScreenTape assay for libraries. Concentrations of amplified cDNA and final libraries were measured using a Qubit 1X dsDNA HS kit (Invitrogen, Waltham, MA) and cDNA amplification and indexing PCR were performed with 12 cycles each. Final libraries were sequenced on an Illumina NovaSeq6000 (Illumina Inc., San Diego, CA).

### Sn-RNA-seq analysis

The mkfastq pipeline in Cell Ranger v3.1 (10X Genomics) was used to convert BCL files into demultiplexed FASTQ files, and the mkref pipeline indexed the *D. melanogaster* reference genome (release 6, GCA_000001215.4 from NCBI Genbank). Alignment was performed using the count pipeline in Cell Ranger v3.1 with the expected cell count set to 5,000. Raw expression counts were imported into Seurat v3.10 in R ^56–58^, normalized using the scTransform ^59^ pipeline with regularized negative binomial regression, and integrated using the SCT method.

Cell clustering was performed using the Louvain algorithm ^60^ with a resolution parameter 1 to optimize community formation. Resolution parameter was selected by iterating between 0.1 to 2.0 to identify stable plateaus that correspond to numbers of cell-type clusters. Dimensionality reduction was achieved using the RunUMAP and FindNeighbors functions with 10 dimensions. Thirty-five cell-type clusters were identified through unsupervised clustering (FindClusters), and annotation of the top 20 markers for each cell type was based on the most frequently occurring identity for cell types using expression scores from BGEE ^61^. Only cell types with an expression score >90 were considered. Annotation of cell type was based on the most frequently occurring identity based on the top 20 marker genes. Clusters 0 and 13 were further subclustered at a resolution of 0.2 to identify the heterogeneous cell identities, resulting in 9 and 4 subclusters, respectively. The subclusters are denoted by appending the subcluster number to the cluster ID, separated by an underscore (“_”).

### Analysis of NaAsO_2_-induced differential expression of genes and genetic networks in the fly brain

Pearson residuals from the scTransform pipeline were used for differential expression analysis. The MAST algorithm in the FindMarkers function was employed to calculate differential expression within clusters after merging clusters with the same identity, incorporating cellular detection rates as a covariate ^62^. *P*-values were adjusted for multiple-hypothesis testing using the Benjamini-Hochberg method and adjusted *P*-values ≤ 0.05 were considered statistically significant.

Gene enrichment analyses were conducted using PAthway, Network, and Gene-set Enrichment Analysis (PANGEA) ^63^. The analysis incorporated GO hierarchy categories (GO Biological Processes, GO Molecular Function, GO Cellular Component) and pathway resources (KEGG ^64^ Pathway *D.mel*, PANTHER ^65^ pathway *D.mel*, REACTOME ^66^ pathway) to identify overrepresented functional categories.

These analyses were performed separately for upregulated and downregulated genes in each cluster and sex, with results further filtered using a Benjamini-Hochberg ^67^ adjusted *P*-value threshold of ≤ 0.05.

The scaled data generated from the sctransform ^59^ pipeline for differentially expressed genes across all clusters in male and female samples were extracted to construct co-expression networks. Pairwise Spearman correlations were computed using these scaled datasets, and the resulting correlation networks were visualized with Cytoscape version 3.7.2 ^68^. To create protein-protein interaction networks, gene IDs were converted to gene symbols using the FlyBase Consortium’s ‘Query-by-symbols/ID’ tool. Interactions between gene products were then calculated by filtering for differentially expressed genes from the known physical interaction database from FlyBase (release v2024_v2) within Cytoscape ^68^. In addition, genes that were connected to at least six differentially expressed genes were computationally recruited and included within the network. The network was subclustered using MCODE plugin and default parameters ^69^. Smaller MCODE clusters with high degree of interconnectivity were merged together. Human orthologs for Drosophila genes were identified using the Drosophila RNAi Screening Center Integrative Ortholog Prediction Tool (DIOPT) v9.0 ^70^. Multiple human orthologs for the same Drosophila gene were resolved by selecting the human gene with the highest DIOPT score. Functional characterization of the subclusters were accomplished using Gene Ontology enrichment analysis and over-representation tests. Only categories with Benjamini-Hochberg adjusted *P*-value ≤ 0.05 for the over-representation tests were considered.

To compare protein-protein interaction networks between Drosophila and humans, the Drosophila network was filtered for nodes with predicted human orthologs (DIOPT ≥ 3). String-DB v11.5 interaction database was filtered for these human orthologs as well as a combined interaction score threshold ≥ 500. The threshold for the combined interaction score was determined by constructing a histogram for distribution of the scores in the filtered network. The two networks were computationally compared using the DyNet plugin within Cytoscape ^71^. We used a Monte Carlo Permutation Procedure to assess the statistical significance of the observed connectivity within the network composed of the shared edges. Briefly, we generated a null distribution of network connectivity indices (average number of neighbors) from randomly sampled subnetworks from the score-filtered protein-protein interaction database, with the same number of genes as the observed network. *P*-value was calculated as the ratio between the rank of the observed connectivity index and the total number of permutations (N = 10,000). Construction of the null distribution using randomly sampled networks and calculation of the connectivity indices were performed using the *igraph* R package v1.4.2 in R.

## Data availability

Raw data for climbing and lifespan measurements are included in the supplementary tables. All single-nuclei RNA sequences data generated in this study have been submitted to the NCBI Gene Expression Omnibus (GEO; https://www.ncbi.nlm.nih.gov/geo/query/acc.cgi?acc=GSE290993) under accession numbers GSE290993, GSM8827136, GSM8827137, GSM8827138 and GSM8827139. R code that was used to perform Seurat-based and Network-based analyses is available at GitHub repositories https://github.com/Anuragbio/As-snRNAseq and https://github.com/vshanka23/mcpp_network_significance respectively.

## Acknowledgements

This project has received funding from the European Union’s Horizon 2020 Research and Innovation program under Grant Agreement No 965406. The work presented in this publication was performed as part of ASPIS. The results and conclusions reflect only the authors’ views, and the European Commission cannot be held responsible for any use that may be made of the information contained therein. AC is supported by a fellowship from the Self Family Foundation. The Genomics and Bioinformatics Cores of the Center for Human Genetics are supported in part by grant 1P20GM139769 from the National Institute of General Medical Sciences of the National Institutes of Health to TFCM and RRHA.

## Author contributions

The project was conceived by AC, TFCM, RRHA. PF dissected brains and generated single nuclei preparations, RAL performed sequencing, EH and KMC performed range finding experiments, BS contributed to the climbing experiments, AC reared the fly lines, exposed them to arsenic and measured survival. AC and VS analysed the data. TFCM and RRHA acquired funding and directed the project. AC, TFCM and RRHA wrote the manuscript.

## Supplementary Figures and Tables

**Fig. S1.** Example of cell cluster annotation. Cluster 11 was annotated as perineurial glial cells based on the BGEE frequency of gene expression for the top 20 marker genes, all of which were exclusively expressed in a single cell type.

**Fig. S2.** Venn diagram that illustrates the differentially expressed genes (DEGs) that are shared and unique between males and females across all clusters.

**Fig. S3.** Venn diagram that illustrates the differentially expressed genes (DEGs) that are shared and unique between males and females for each cluster separately.

**Fig. S4.** Heatmap showing the expression patterns of 412 genes common to both males and females (Blue: downregulated, Red: upregulated).

**Fig. S5:** Histogram of the null distribution with kernel density estimation overlay calculated for network connectivity indices from the randomly sampled subnetworks.

**Supplementary Table S1. Raw data for lifespan.**

**Supplementary Table S2. ANOVA for survival and climbing assay.**

**Supplementary Table S3. Raw data for climbing assay.**

**Supplementary Table S4. Summary of sequencing statistics.**

**Supplementary Table S5. Marker genes used to annotate cell clusters, including subclusters of 0 and 13.**

**Supplementary Table S6. Differentially expressed genes (DEGs) identified in each cluster for males and females**.

**Supplementary Table S7. 26 DEGs in males and 10 DEGs in females exhibiting distinct expression profiles in multiple specific cell types.**

**Supplementary Table S8. Gene enrichment analysis of differentially expressed genes (DEGs) for males and females.** Analyses were performed using PANGEA: GO Biological Processes, PANTHER Pathways (*Drosophila melanogaster*), GO Cellular Components, REACTOME Pathways, GO Molecular Functions, and KEGG Pathways (*Drosophila melanogaster*).

**Supplementary Table S9. Global gene enrichment analysis of differentially expressed genes (DEGs) for all clusters of males and females.**

**Supplementary Table S10. The frequency of each differentially expressed gene across multiple cell types and their specific cluster identities in Drosophila.**

