## Supplementary figures and images for "Arsenic Toxicity in the Drosophila Brain at Single Cell Resolution"

### Supplemental Figure 1

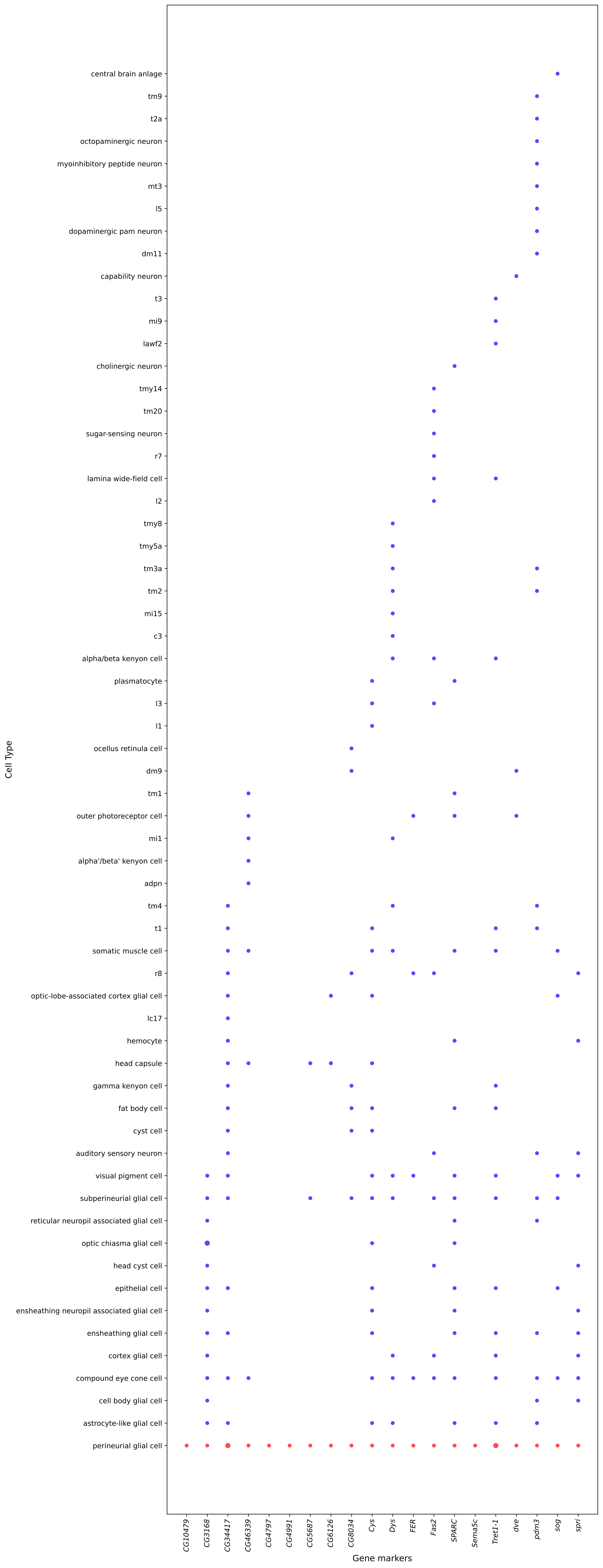

### Supplemental Figure 2

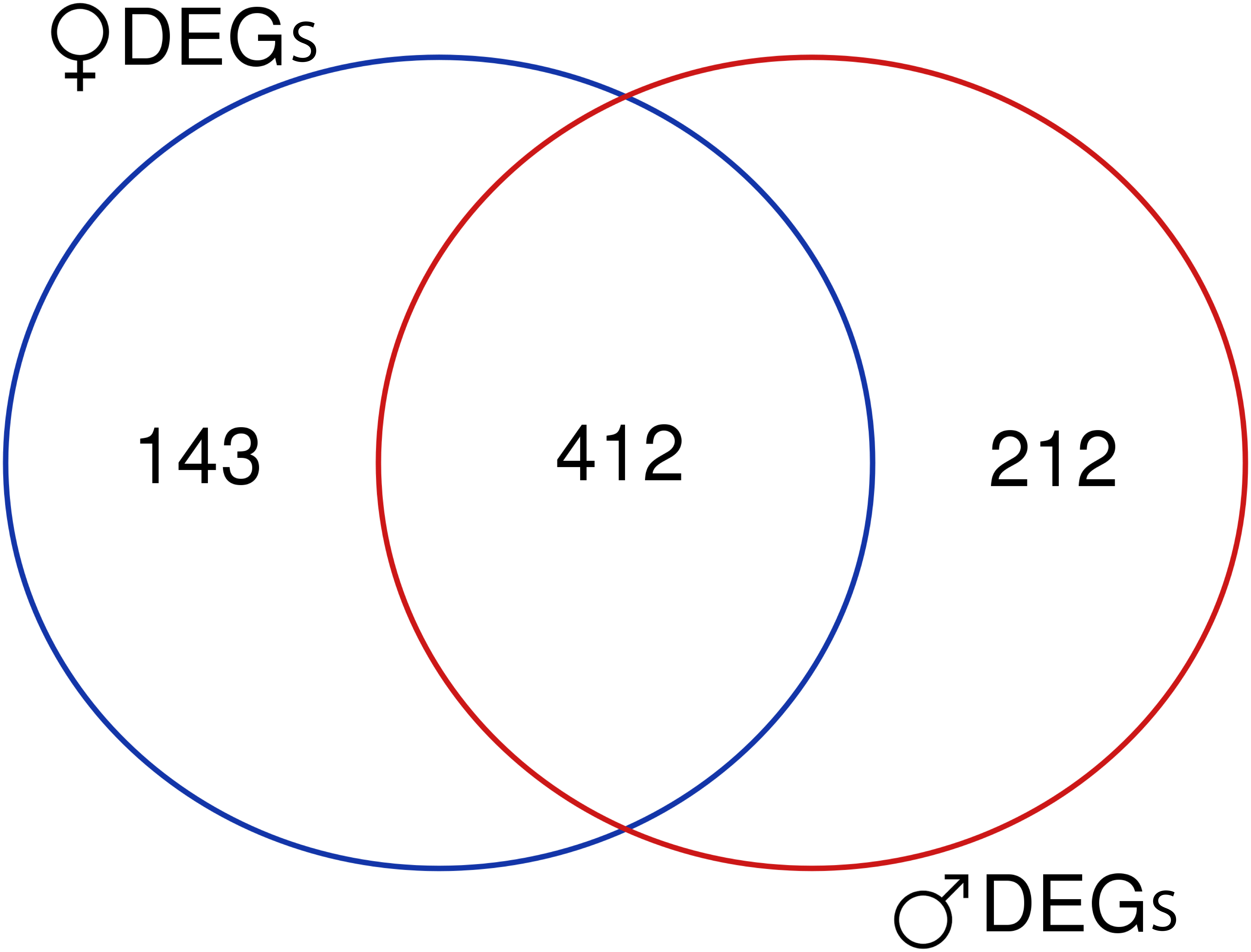

### Supplemental Figure 3

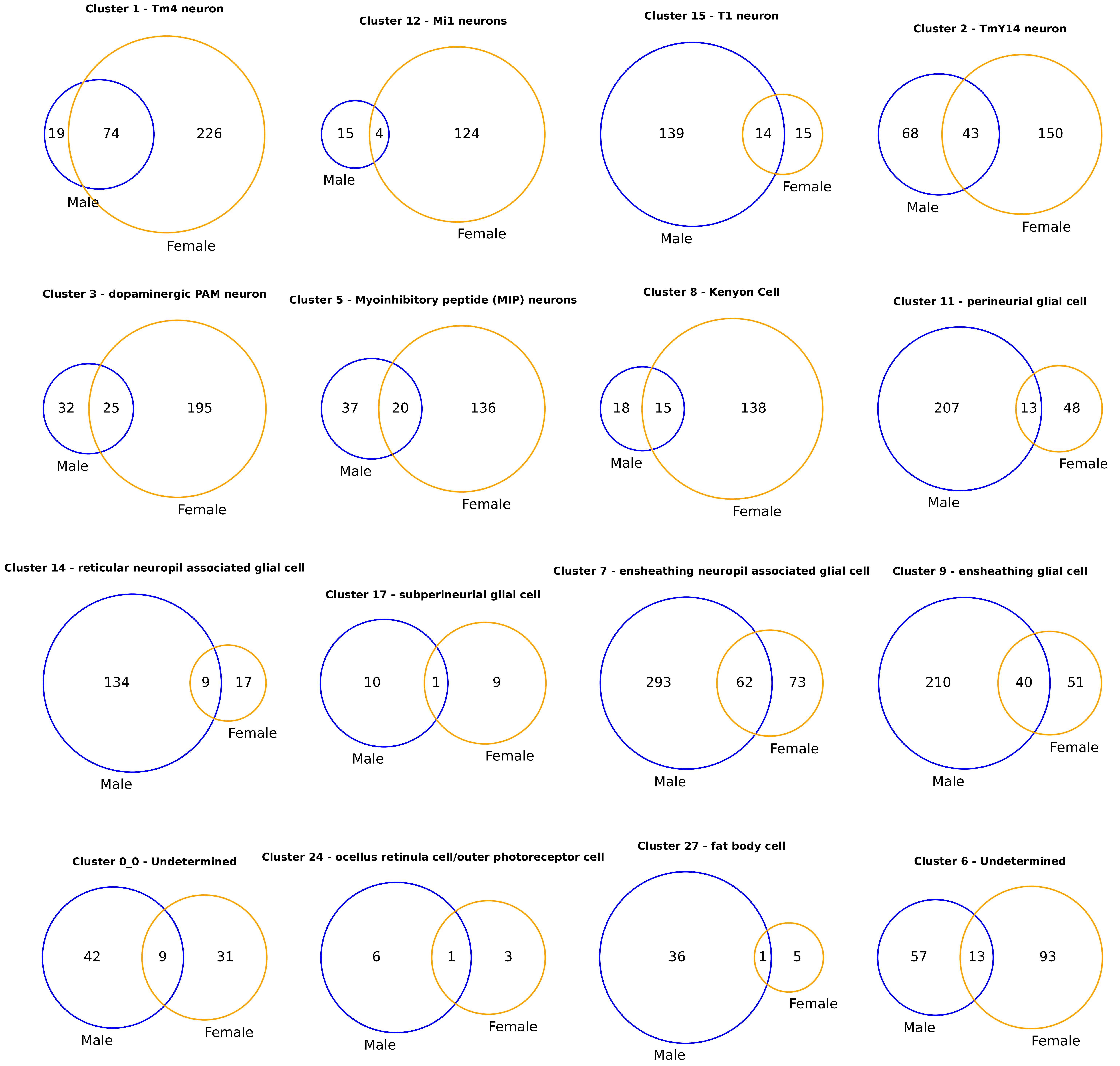

### Supplemental Figure 4

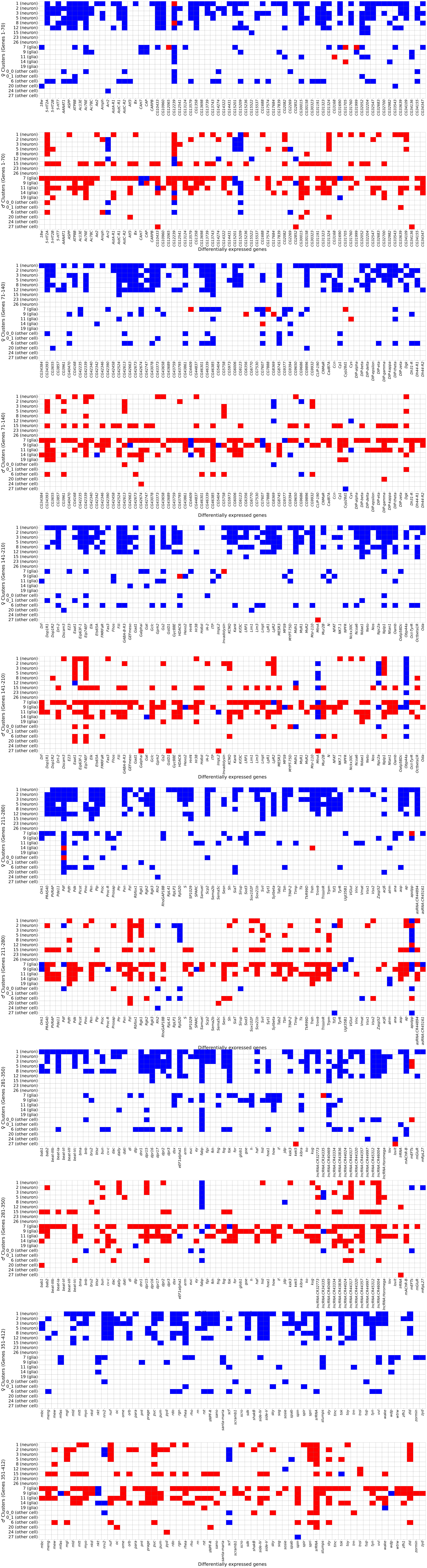

### Supplemental Figure 5

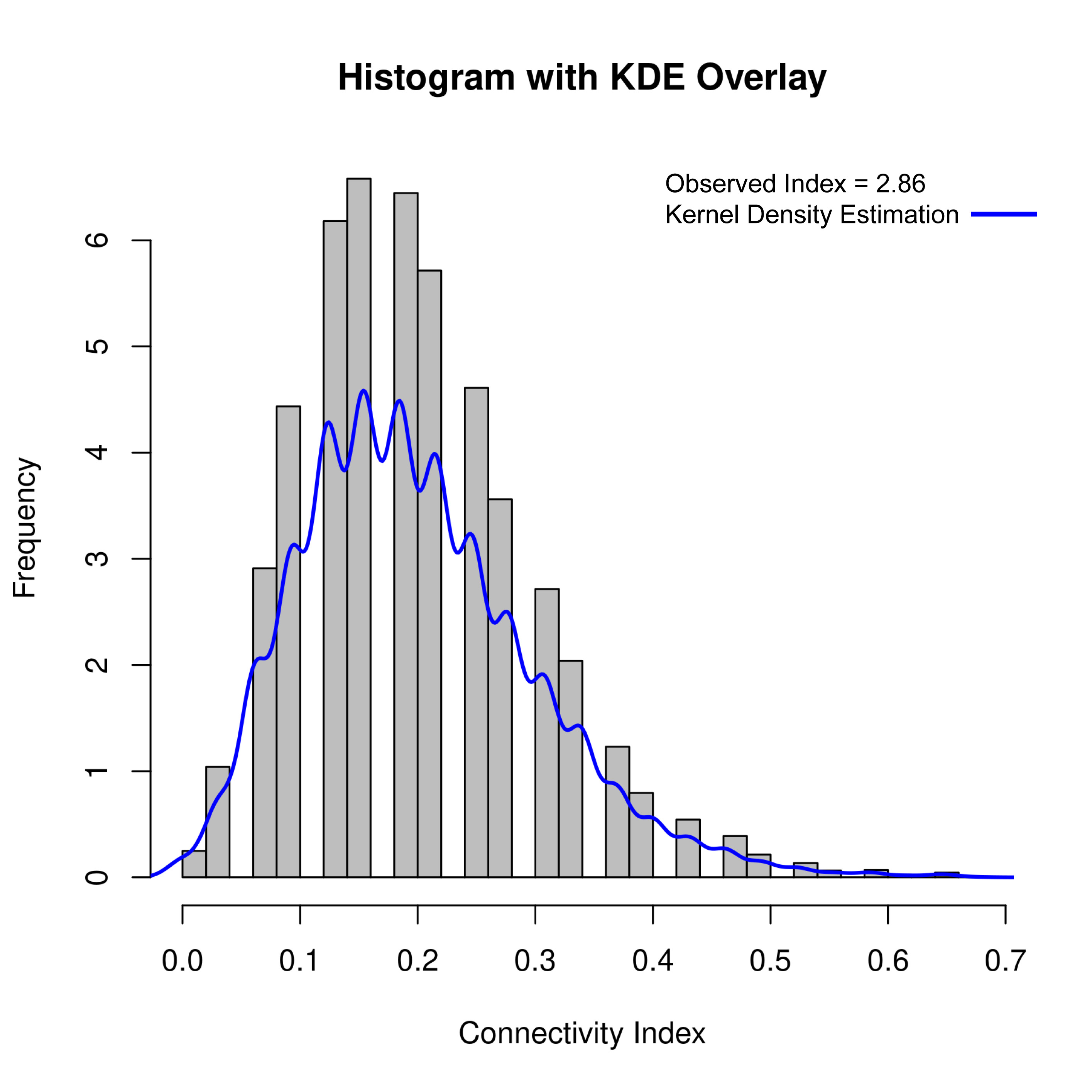
